# The anticoagulant nafamostat potently inhibits SARS-CoV-2 infection *in vitro*: an existing drug with multiple possible therapeutic effects

**DOI:** 10.1101/2020.04.22.054981

**Authors:** Mizuki Yamamoto, Maki Kiso, Yuko Sakai-Tagawa, Kiyoko Iwatsuki-Horimoto, Masaki Imai, Makoto Takeda, Noriko Kinoshita, Norio Ohmagari, Jin Gohda, Kentaro Semba, Zene Matsuda, Yasushi Kawaguchi, Yoshihiro Kawaoka, Jun-ichiro Inoue

**Affiliations:** Research Center for Asian Infectious Diseases, Institute of Medical Science, University of Tokyo, Tokyo 108-8639, Japan; Division of Virology, Department of Microbiology and Immunology, Institute of Medical Science, University of Tokyo, Tokyo 108-8639, Japan; Department of Virology 3, National Institute of Infectious Diseases, Musashimurayama, Tokyo 208-0011, Japan; Disease Control and Prevention Center, National Center for Global Health and Medicine, Tokyo 162-8655, Japan; Department of Life Science and Medical Bio-Science, Waseda University, Shinjuku-ku, Tokyo, 162-8480, Japan; Division of Molecular Virology, Department of Microbiology and Immunology, The Institute of Medical Science, The University of Tokyo, Tokyo 108-8639, Japan; Influenza Research Institute, Department of Pathobiological Sciences, School of Veterinary Medicine, University of Wisconsin-Madison, WI 53711, USA; Department of Special Pathogens, International Research Center for Infectious Diseases, Institute of Medical Science, University of Tokyo, Tokyo 108-8639, Japan; Senior Professor Office, University of Tokyo, Tokyo 113-0033, Japan

**Author notes:** Correspondence: Jun-ichiro Inoue.

## Abstract

Although infection by SARS-CoV-2, the causative agent of COVID-19, is spreading rapidly worldwide, no drug has been shown to be sufficiently effective for treating COVID-19. We previously found that nafamostat mesylate, an existing drug used for disseminated intravascular coagulation (DIC), effectively blocked MERS-CoV S protein-initiated cell fusion by targeting TMPRSS2, and inhibited MERS-CoV infection of human lung epithelium-derived Calu-3 cells. Here we established a quantitative fusion assay dependent on SARS-CoV-2 S protein, ACE2 and TMPRSS2, and found that nafamostat mesylate potently inhibited the fusion while camostat mesylate was about 10-fold less active. Furthermore, nafamostat mesylate blocked SARS-CoV-2 infection of Calu-3 cells with an EC_50_ around 10 nM, which is below its average blood concentration after intravenous administration through continuous infusion. These findings, together with accumulated clinical data regarding its safety, make nafamostat a likely candidate drug to treat COVID-19.

---

Infection by severe acute respiratory syndrome coronavirus 2 (SARS-CoV-2), the causative agent of coronavirus pneumonia disease (COVID-19), is spreading rapidly worldwide^1^. As yet, no drug has been shown to be sufficiently effective for treating COVID-19. Therefore, drug repurposing offers potentially the quickest path toward disease treatment.

The genomic RNA of coronaviruses is surrounded by an envelope^2^. Initiation of viral entry requires two steps: 1) the Spike (S) protein in the viral envelope, binds to its receptor present in the plasma membrane after S protein is cleaved into S1 and S2 proteins by the cellular protease. SARS-CoV and SARS-CoV-2 use angiotensin converting enzyme 2 (ACE2), while MERS-CoV uses CD26 as a receptor; 2) S2 is cleaved by a cell surface transmembrane protease, serine 2 (TMPRSS2), so-called priming, which exposes the fusion peptide in S2 protein, allowing it to attach to the plasma membrane resulting in the fusion between envelope and the plasma membrane (envelope fusion). This fusion allows the viral RNA to enter the cytoplasm, where it replicates. TMPRSS2-knockout resulted in reduced spread of SARS-CoV and MERS-CoV in the airways accompanied by reduced severity of lung pathology in a mouse model^3^. Therefore, TMPRSS2 is likely crucial for SARS-CoV-2 spreading and disease development *in vivo*, and targeting it either by inhibiting its priming activity or suppressing its expression is likely to be an effective strategy to cure COVID-19.

We have previously reported that nafamostat mesylate, an existing Japanese drug used for acute pancreatitis and disseminated intravascular coagulation (DIC) with enhanced fibrinolysis, effectively inhibits MERS-CoV S protein-initiated membrane fusion by targeting TMPRSS2 priming activity^4^. We did this using the cell fusion assay monitored by the Dual Split Protein (DSP) reporter to screen the FDA-approved drug library (Fig. 1a). Nafamostat mesylate potently inhibited MERS-CoV infection of lung epithelium-derived Calu-3 cells.

**Fig. 1.**
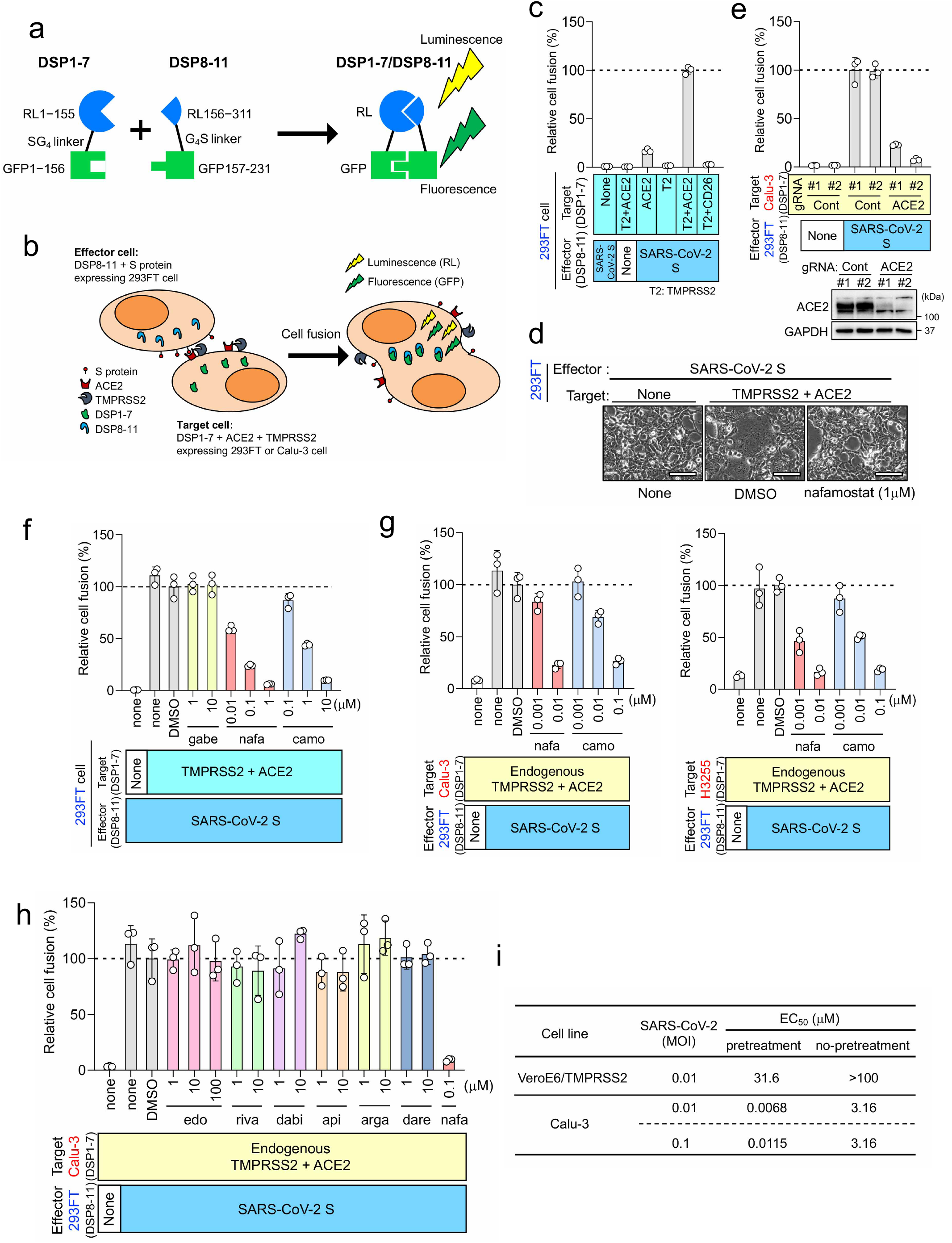
Nafamostat mesylate potently inhibits TMPRSS2-dependent SARS-CoV-2 entry into lung epithelium-derived Calu-3 cells. **a** DSP1-7 has the structure RL_1–155_-Ser-Gly-Gly-Gly-Gly-GFP_1–156_. DSP8-11 has the structure Met-RL_156–311_-Gly-Gly-Gly-Gly-Ser-GFP_157–231_. DSP1-7 and DSP8-11 reassociate efficiently, resulting in reconstitution of functional RL and GFP to generate luminescent and fluorescent signals, respectively. **b** Effector cells (293FT cells expressing DSP8-11 and S protein) and target cells (293FT or Calu-3 cells expressing DSP1-7, ACE2 and TMPRSS2) were co-cultured. Both GFP (fluorescence) and RL (luminescence) signals were generated following DSP1-7 and DSP8-11 reassociation upon cell fusion. **c** Different combinations of the effector and target cells were cocultured, and the resulting RL activity was measured. Relative cell-fusion values were calculated by normalizing the RL activity of each co-culture to that of the co-culture of cells expressing S protein with those expressing both receptor and TMPRSS2, which was set to 100%. **d** Phase contrast images of SARS-CoV-2 S protein-initiated-cell fusion. Scale bars, 100 μm. **e** The fusion assay using wild type and ACE2-kockout Calu-3 cells. **f** Three clinically used pancreatitis and/or anticoagulant drugs were evaluated by the DSP assay for their effects on SARS-CoV-2 S-initiated membrane fusion. Relative cell-fusion value was calculated by normalizing the RL activity for each co-culture to that of the co-culture with DMSO alone, which was set to 100%. gabe: gabexate mesylate, nafa: nafamostat mesylate, camo: camostat mesylate. **g** The DSP assay using Calu3 (left) or H3255 (right) cells as target cells. DSP1-7 was constitutively expressed in Calu3 and H3255 cells. **h** The DSP assay using Calu-3 cells was performed in the presence of various anticoagulants. edo, edoxaban; riva, rivaroxaban; dabi, dabigatran; api, apixaban; arga, argatroban; dare, darexaban. **i** VeroE6/TMPRSS2 or Calu-3 cells were either untreated or pretreated with nafamostat mesylate (1-10^5^ nM) for 1 h. SARS-CoV-2 was then added followed by 30 min incubation, and the culture medium was changed to fresh medium with the same concentration of nafamostat mesylate. Three or 5 days after infection, EC_50_ was determined. Experiments were done in quadruplicate.

Based on our previous work^4^ together with others’^5^, we established an experimental system monitoring SARS-CoV-2 S protein-initiated membrane fusion, and tested ACE2 and TMPRSS2 dependence in 293FT cells, which do not express ACE2 and TMPRSS2 (Fig. 1b-d). Fusion did not occur when target cells lacked either ACE2 or TMPRSS2 (Fig. 1c), indicating that the cell fusion clearly depends on ACE2 and TMPRSS2 as previously reported for MERS-CoV S protein-initiated fusion, which depends on CD26 and TMPRSS2^4^ (Supplementary information, Fig. S1a). Furthermore, when ACE2-kockout Calu-3 cells, endogenously expressing ACE2 and TMPRSS2, were used as target cells, fusion scarcely occurred, suggesting that no other functional receptor exists in some types of lung epithelial cells (Fig. 1e).

We then tested the activity of three existing Japanese drugs used for pancreatitis and/or DIC (nafamostat mesylate, camostat mesylate and gabexate mesylate) with inhibitory activity against serine proteases to inhibit SARS-CoV-2 S protein-initiated fusion (Fig. 1f). Luciferase activities derived from cells carrying the preformed DSP1-7/DSP8-11 reporter complex were not affected by the drugs (Supplementary information, Fig. S1b), indicating that the suppression of luciferase activities reflects the inhibition of fusion by drugs. Nafamostat mesylate showed greatest activity, camostat mesylate was 10-fold less active, and gabexate mesylate was inactive within the range of concentrations tested (10 nM-10 μM) (Fig. 1d, f), which mirrors drug sensitivity profiles to MERS-CoV S protein-initiated fusion^4^ (Supplementary information, Fig. S1c).

We next used lung epithelium-derived Calu-3 and H3255 cells as target cells, because they endogenously express ACE2 and TMPRSS2, and might better reflect physiological conditions than 293FT kidney cells ectopically expressing ACE2 and TMPRSS2. Nafamostat mesylate inhibited SARS-CoV-2 S protein-initiated membrane fusion in the range of 1-10 nM while camostat mesylate required 10 −100 nM to achieve a similar extent of inhibition (Fig. 1g). A similar inhibition profile was observed when MERS-CoV S protein-initiated fusion was analyzed^4^ (Supplementary information, Fig. S1d). Therefore, nafamostat mesylate was 10-fold more potent than camostat mesylate. Furthermore, drug concentrations required for fusion inhibition in lung epithelial cells were 10-fold lower than those required for inhibition using 293FT cells (Fig. 1f, g).

Various computational approaches have recently been applied to find existing medications targeting TMPRSS2^6,7^. Consistent with our results, these studies ranked nafamostat mesylate significantly higher than camostat mesylate. It should be noted that several world-wide commercially available anticoagulants, which are serine protease inhibitors targeting Factor Xa or Thrombin, are listed. Therefore, we checked whether these anticoagulants affect SARS-CoV-2 S protein-initiated membrane fusion. Contrary to expectations, they all failed to inhibit membrane fusion (Fig. 1h).

Our DSP assays identified nafamostat mesylate as a potent inhibitor of SARS-CoV-2 S-initiated membrane fusion (Fig. 1f, g). This inhibition was most likely a result of inhibition of TMPRSS2 on the plasma membrane^2^. SARS-CoV-2 infects Calu-3 cells primarily via the TMPRSS2-dependent plasma membrane pathway^4,5,8^, while it infects VeroE6/TMPRSS2 cells via the TMPRSS2-independent endocytotic pathway in addition to the plasma membrane pathway^9^. We then proceeded to evaluate the effects of nafamostat mesylate on actual SARS-CoV-2 infection in both types of cells with or without pretreatment with nafamostat mesylate. Based on the appearance of cytopathic effect (CPE), EC_50_ values for Calu-3 cells were 6.8-11.5 nM (with pretreatment) and 3.16 μM (without pretreatment) while those for VeroE6/TMPRSS2 cells were 31.6 μM (with pretreatment) and >100 μM (without pretreatment) (Fig. 1i). In both cell types, EC_50_ values were much smaller when cells were pretreated with nafamostat mesylate, implying that nafamostat mesylate inhibits SARS-CoV-2 entry. More importantly, EC_50_ values for Calu-3 cells with the pretreatment were around 10 nM, similar to our previous findings with MERS-CoV infection^4^. This extremely high sensitivity to nafamostat mesylate may be because the TMPRSS2-dependent infection pathway is dominant in lung epithelium-derived Calu-3 cells^4,5,8^. Given that camostat mesylate-mediated inhibition of TMPRSS2 significantly reduced SARS-CoV-2 infection of primary human airway epithelial cells^5^ and that blood concentrations of nafamostat mesylate were maintained at 30–240 nM when it was administered intravenously through continuous infusion according to the standard protocol for DIC patients^10^, nafamostat mesylate could be used to treat COVID-19. In contrast, significantly higher doses of nafamostat mesylate are required to block SARS-CoV-2 infection of monkey kidney VeroE6/TMPRSS2 cells as reported previously^11^, which is probably due to the significant contribution of the TMPRSS2-independent endocytotic infection pathway. Because a pharmacokinetics study using rats revealed the maximum concentration of intact nafamostat mesylate in the lung after infusion to be about 60-fold higher in comparison with the maximum blood concentration^12^, such an accumulation may partially suppress SARS-CoV-2 infection of low-responder cells like VeroE6/TMPRSS2 cells. Therefore, identification and characterization of cells that play a key role in virus spread and disease development is required.

Our findings clearly indicate that nafamostat mesylate, the most effective TMPRSS2 inhibitor so far reported, potently inhibits SARS-CoV-2 infection *in vitro*. Hoffmann et al. have recently reported that Nafamostat mesylate blocks activation of SARS-CoV-2^13^. However here we clearly demonstrate that nafamostat mesylate blocks membrane fusion step of the virus entry and its activity to block SARS-CoV-2 infection is cell type dependent. These findings are crucial for developing therapeutic strategies. It has been reported recently that abnormal coagulation with elevated concentrations of D-dimer, characteristic of DIC with enhanced fibrinolysis, may influence the prognosis of COVID-19^14,15^. Furthermore, in a murine asthma model, nafamostat mesylate attenuates respiratory inflammation by blocking activation of NF-κB, a critical transcription factor for inflammatory cytokine production^16^. Therefore, nafamostat mesylate is expected to have multiple therapeutic effects. Because nafamostat mesylate has been prescribed in Japan for many years and adequate clinical data regarding safety have accumulated, we suggest that it should be evaluated in COVID-19 patients by itself or in combination with other antiviral drugs that target separate processes needed for virus production.

## Material and Methods

### Protease inhibitors and anti-coagulants

Nafamostat mesylate (Tokyo Chemical Industry, Tokyo, Japan), camostat mesylate (Wako, Tokyo, Japan), gabexate mesylate (Tokyo Chemical Industry, Tokyo, Japan), edoxaban, apixaban, rivaroxaban, dabigatran (Selleck Chemicals, Houston, TX, USA), argatroban (Tokyo Chemical Industry, Tokyo, Japan) and darexaban (Santa Cruz Biotechnology, SantaCruz, CA, USA)

### Cell lines and transient transfection

HEK293FT is an immortalized cell line derived from human fetal kidney. A pair of previously described 293FT-based reporter cell lines that constitutively express individual split reporters (DSP1-7 and DSP8-11 proteins)^17^ were used in this study and maintained in Dulbecco’s modified Eagle’s medium (DMEM) containing 10% fetal bovine serum (FBS) and 1 μg/ml puromycin. For establishment of stable cell lines expressing the S protein of SARS-CoV-2, or MERS-CoV, recombinant pseudotype lentiviruses were produced using HEK293T cells with lentiviral transfer plasmid expressing S protein, psPAX2 packaging plasmid and vesicular stomatitis virus (VSV)-G-expressing plasmid. For establishment of stable cell lines expressing ACE2 or CD26, and TMPRSS2, recombinant pseudotype retroviruses expressing one of these proteins were produced using plat-E cells with a VSV-G-expressing plasmid^18^. 293FT-derived reporter cells infected with pseudotype viruses were selected with 1 μg/ml puromycin, 10 μg/ml blasticidin, and 300 μg/ml hygromycin for at least 1 week. These bulk selected cells were used to perform fusion assays. Calu-3 cells (ATCC HTB-55) and H3255 cells (CVCL_6831), lung epithelial cell-derived immortalized cells established from human lung cancer, were used as target cells for the fusion and viral infection assays. For establishment of ACE2-knockout Calu-3 cells, lentiviruses were produced by transfecting lentiCRISPRv2 vector (#52961 Addgene, Watertown, MA, USA) with the following gRNA sequences. The gRNA sequences used were 5’-GCT TTC ACG GAG GTT CGA CG and 5’-ATG TTG CAG TTC GGC TCG AT for control, and 5’-ATG AGC ACC ATC TAC AGT AC and 5’-TGC TGC TCA GTC CAC CAT TG for ACE2-knockout. Pooled Calu-3 cells infected with pseudotype viruses were selected with 1 μg/ml puromycin.

### Construction of expression vectors

Expression vectors for CD26 and TMPRSS2 were described previously^4^. A synthetic DNA corresponding to the S gene of SARS-CoV-2 (NC_045512.2) was generated by Taihe Gene (Beijing, China). For construction of expression vectors for ACE2, the ACE2 gene was cloned into a pMXs-internal ribosome entry site (IRES)-blasticidin retroviral vector^2^. For construction of expression vectors for S protein of SARS-CoV-2 and MERS-CoV, the coding regions were cloned into a lentiviral transfer plasmid (CD500B-1, SBI, Palo Alto, CA, USA).

### DSP assay to monitor membrane fusion

For the DSP assay using 293FT cells, effector cells expressing S protein with DSP8-11 and target cells expressing CD26 or ACE2, and TMPRSS2 with DSP1-7 were seeded in 12-well cell culture plates (2 x 10^5^ cells/500 μl) one day before the assay. Two hours before the DSP assay, cells were treated with 6 μM EnduRen (Promega, Madison, WI, USA), a substrate for Renilla luciferase, to activate EnduRen. One μl of each protease inhibitor or anticoagulant dissolved in dimethyl sulfoxide (DMSO) was added to the 384-well plates (Greiner Bioscience, Frickenhausen, Germany). Next, 50 μl of each single cell suspension (effector and target cells) was added to the wells using a Multidrop dispenser (Thermo Scientific, Waltham, MA, USA). After incubation at 37°C for 4 h, the RL activity was measured using a Centro xS960 luminometer (Berthold, Germany). For the DSP assay using Calu-3 or H3255 cells, target cells were seeded in 384-well plates (2 x 10^4^ cells/50 μl) one day before the assay. Two hours before the DSP assay, cells were treated with 6 μM EnduRen. One μl of each protease inhibitor or anticoagulant dissolved in DMSO was added to the 384-well plates with 9 μl of culture medium. Next, 40 μl of single cell suspension (effector cells) was added to the wells using a Multidrop dispenser.

### Western blotting

Western blot analysis was performed as described previously^19^. The primary antibodies used were rabbit anti-ACE2 (1:1000, ab15348 Abcam, Cambridge, MA, USA) and anti-GAPDH (1:1000, sc-25778 Santa Cruz Biotechnology, Dallas, TX, USA). HRP-linked donkey anti-rabbit IgG (NA934, GE Healthcare, Piscataway, NJ, USA).

### Isolation of SARS-CoV-2

VeroE6 (ATCC CRL-1586) cells were maintained in Eagle’s minimal essential media (MEM) containing 10% FBS. The cells were incubated at 37 °C with 5% CO2, and regularly tested for mycoplasma contamination by using PCR and were confirmed to be mycoplasma-free. Respiratory swabs were obtained from a patient with laboratory-confirmed COVID-19, who was hospitalized at the Center Hospital of the National Center for Global Health and Medicine, Tokyo, Japan. The swabs were submitted to the Division of Virology, Department of Microbiology and Immunology, Institute of Medical Science, the University of Tokyo for virus isolation by inoculating with VeroE6 cells. The research protocol was approved by the Research Ethics Review Committee of the Institute of Medical Science of the University of Tokyo. SARS-CoV-2 viruses were propagated in VeroE6 cells in Opti-MEM I (Invitrogen) containing 0.3% bovine serum albumin (BSA) and 1 μg of L-1-Tosylamide-2-phenylethyl chloromethyl ketone (TPCK)-trypsin/ml at 37 °C. All experiments with SARS-CoV-2 viruses were performed in enhanced biosafety level 3 (BSL3) containment laboratories at the University of Tokyo, which are approved for such use by the Ministry of Agriculture, Forestry, and Fisheries, Japan.

### SARS-CoV-2 infection assay

Calu-3 (ATCC HTB-55) cells were maintained in MEM containing 10% FBS. VeroE6/TMPRSS2^20^ (JCRB 1819) cells were propagated in the presence of 1 mg/ml geneticin (G418; Invivogen) and 5 μg/ml plasmocin prophylactic (Invivogen) in DMEM containing 10% FBS. All cells were incubated at 37 °C with 5% CO2, and regularly tested for mycoplasma contamination by using PCR and were confirmed to be mycoplasma-free. The target VeroE6/TMPRSS2 or Calu-3 cells were untreated or pretreated with nafamostat mesylate (1-10^5^ nM, 4 wells for each dose) for 1 h, and SARS-CoV-2 was then added at a multiplicity of infection (MOI) of 0.01 or 0.1. The viruses and cells were incubated for 30 min for viral entry, and the culture medium was changed to fresh medium with the same concentration of nafamostat mesylate. Three or 5 days after infection, EC_50_ was determined based on the appearance of visually detectable cytopathic effect (CPE). Experiments were done in quadruplicate.

## Acknowledgements

We thank Shin-ichiro Hattori for clinical specimens, Soko Takahashi, Kinuyo Miyazaki and Kiyomi Nakagawa for their secretarial assistance. This work was supported in part by grants-in-aid from the Japanese Society for the Promotion of Science (16H06575 to JI, 18K15235 to MY), a Program of Japan Initiative for Global Research Network on Infectious Diseases (JGRID) from AMED (JP18fm0108006 to MY, YaK, Yok, ZM, and JI), NIAID-funded Center for Research on Influenza Pathogenesis (CRIP; HHSN272201400008C to YoK) and a Research Program on Emerging and Re-emerging Infectious Diseases from AMED (19fk0108113 to YoK).

## Author contributions

M.Y., M.K., J.G., Z.M., Ya.K., Yo.K. and J.I. conceived and designed the experiments. M.Y., M.K., Y.S.-T., K.I.-H., M.I., M.T., N.K., N.O. and J.G. performed the experiments. M.Y., M.K., M.I., J.G., Z.M., K.S., Ya.K., Yo.K. and J.I. analyzed the data. M.Y. and J.I. wrote the paper.

## Additional Information

### Supplementary Information

#### Competing Interests

The authors declare no competing interests.

## Notes

### Competing Interest Statement

The authors have declared no competing interest.

