## Supplementary Fig. S1 for "The anticoagulant nafamostat potently inhibits SARS-CoV-2 infection *in vitro*: an existing drug with multiple possible therapeutic effects"

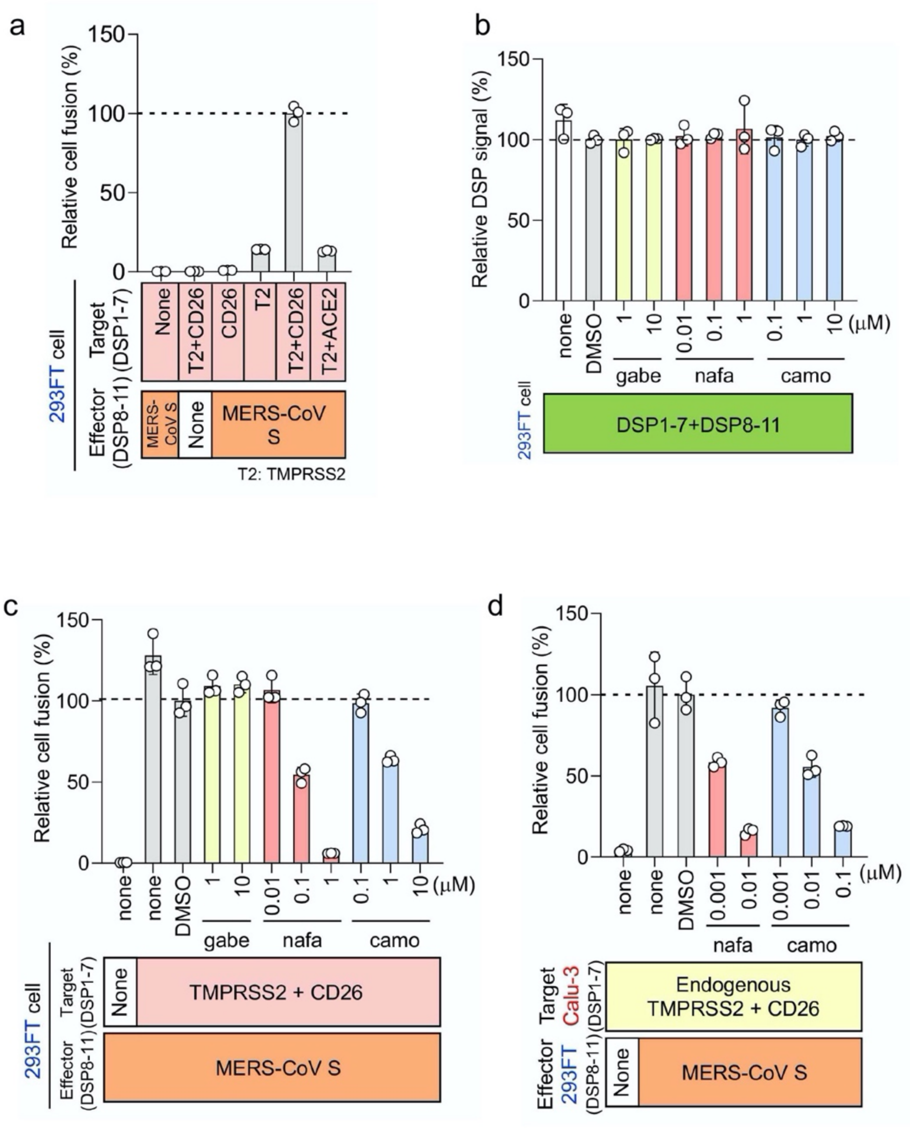

**Fig S1. Nafamostat mesylate potently inhibits the TMPRSS2- and CD26-dependent MERS-CoV S protein-initiated membrane fusion.**

**a** Quantitative assay for the TMPRSS2- and CD26-dependent MERS-CoV S protein-initiated membrane fusion. The effector 293FT cells expressing DSP8-11 were transduced with MERS-CoV S, and the target 293FT cells expressing DSP1-7 were transduced with CD26 and TMPRSS2 separately or simultaneously. Different combinations of these effector and target cells were cocultured, and the resulting RL activity was measured. Relative cell-fusion value was calculated by normalizing the RL activity for each co-culture to the RL activity of the co-culture of effector cells expressing S protein and target cells expressing both receptor and TMPRSS2, which was set to 100%.
**b** The effect of nafamostat mesylate, camostat mesylate and gabexate mesylate on RL measurement. Each drug was added to cells co-expressing DSP1-7 and DSP8-11 to evaluate any direct inhibitory effects on RL. The relative DSP signal is indicated in the vertical axis by setting the control value with DSP alone as 100%.
**c** Nafamostat mesylate potently inhibits MERS-CoV S protein-initiated membrane fusion of 293FT cells. Nafamostat mesylate, camostat mesylate and gabexate mesylate, were evaluated by the DSP assay for their effects on MERS-CoV S-initiated membrane fusion. The effect of each drug on the coculture fusion assay using DSP as a reporter. Drugs were tested at different concentrations, and the additional proteins transduced into the effector and target cells are indicated below the graph. Relative cell-fusion value was calculated by normalizing the RL activity for each co-culture to that of the co-culture effector cells expressing S protein and target cells expressing both receptor and TMPRSS2 with DMSO alone, which was set to 100%. gabe, gabexate mesylate; nafa, nafamostat mesylate; camo, camostat mesylate.
**d.** Lung epithelium-derived Calu-3 cells were more sensitive to nafamostat mesylate and camostat mesylate compared with 293FT cells in inhibiting MERS-CoV S protein-initiated membrane fusion. The DSP assay system using Calu-3 cells as target cells. DSP1-7 was constitutively expressed in Calu-3 cells. Using 293FT-derived effector cells expressing DSP8-11 and MERS-CoV S protein, the DSP assay was performed in the presence of nafamostat mesylate or camostat mesylate. Relative cell-fusion value was calculated by normalizing the RL activity for each co-culture to the RL activity of the co-culture of cells expressing S protein with Calu-3 cells with DMSO alone, which was set to 100%.
